# The Developing Human Connectome Project: Fetal Data Release

**DOI:** 10.64898/2026.11.25.768215

**Authors:** A. David Edwards, Samy Abo Seada, Amir Alansary, Jesper Andersson, Tomoki Arichi, Dafnis Batalle, Arnaud Boutillon, Jelena Bozek, Eleanor Braithwaite, Olivia Carney, Andrew Chew, Daan Christiaens, Raymond Chung, Naomi Clarke, Kathleen Colford, Lucilio Cordero-Grande, Serena J. Counsell, Harriet Cullen, John Cupitt, Charles Curtis, Alice Davidson, Maria Deprez, Mathieu Deprez, Louise Dillon, Konstantina Dimitrakopoulou, Eugene Duff, Shona Falconer, Seyedeh-Rezvan Farahibozorg, Mona Farhangi, Sean P Fitzgibbon, Jianliang Gao, Andreia Sofia Oliveira Gaspar, Terrenjit Gill, Irina Grigorescu, Yourong Guo, Nicholas Harper, Emer J Hughes, Jana Hutter, Mark Jenkinson, Saad Jbabdi, Emily Jones, Sadia Joulhash, Vyacheslav Karolis, Vanessa Kyriakopoulou, Gregor Lenz, Kaili Liang, Antonios Makropoulos, Shaihan Malik, Luke Mason, Rebecca Morris, Filippo Mortari, Chiara Nosarti, Rita G Nunes, Jonathan O’Muircheartaigh, Hamel Patel, Jonathan Passerat-Palmbach, Maximilian Pietsch, Denis Prokopenko, Anthony N.Price, Cristina Ramirez Piriz, Emma C. Robinson, Daniel Rueckert, Mary A. Rutherford, Andreas Schuh, Nitharshini Sivalingam, Stephen M Smith, Helena Sousa, Rui Pedro Azeredo Gomes Teixeira, Tencho Tenev, Jacques-Donald Tournier, Alena Uus, Krishnika Vetrivel, Logan Z.J. Williams, Siân Wilson, Robert Wright, Wenxuan Wu, Jiaxin Xiao, Charlie Yuli Zhang, Joseph V. Hajnal

**Author notes:** Corresponding author: Professor JV Hajnal, Centre for the Developing Brain, School of Biomedical Engineering and Imaging Sciences, King’s College London, London, SE1 7EH, UK.

## Abstract

The Developing Human Connectome Project (dHCP) represents a large open science resource designed to enable detailed investigations of typical and atypical brain development across the perinatal period. It consists of 1228 multimodal magnetic resonance image examinations of the fetal and/or neonatal brain from 1173 participants, together with collateral demographic, clinical, family, neurocognitive and genomic data. All subjects were studied *in utero* and/or soon after birth with a single MRI scanner using specially developed scanning sequences with novel motion-tolerant imaging methods and optimised for the changing properties of the developing brain. The neonatal data has been described and released separately. The fetal data release consists of multimodal data from 273 subjects scanned between 20 + 6 and 38+2 weeks (+days) of gestation, with some fetuses imaged more than once to provide longitudinal data creating 297 datasets in total. The data include: high resolution and motion corrected T1 and T2 weighted anatomical images; high temporal resolution functional and multi-shell high angular resolution diffusion MRI; and multimodal age-resolved atlases; together with demographic, family and clinical data, genomic and epigenomic data and follow-up neurodevelopmental assessment at around 18 months. This paper describes the dHCP fetal image acquisition, reconstruction and processing protocols, summarizes the available data, and provides information on how it can be accessed.

## Introduction

In recent years, dramatic new insight into the human brain’s architecture of macroscopic connections have been gained through marked advances in Magnetic Resonance Imaging (MRI) data acquisition, processing and analysis in major open science initiatives such as the Human Connectome Project (Craddock et al., 2013; Van Essen et al., 2013). This has included knowledge not only about the framework of white matter pathways that make up the structural connections (Zalesky & Di Biase, 2026), but also the spatial topography of correlated neural connectivity representing the brain’s functional connections (Cosío-Guirado et al., 2024), and its coordinated anatomical development (Alexander-Bloch et al., 2013).

A compelling and crucial challenge is knowledge of how the brain’s connectome is first established in the time leading up to and around birth. Across this time, dramatic changes are known to occur as the brain not only rapidly grows in size but also in the composition and complexity of its tissue as neural progenitor cells form, differentiate, and migrate to form key structures including the emerging white matter pathways and the immature cortex (Molnár et al., 2019). It is increasingly appreciated that this period is crucial for lifelong brain health, with multiple lines of evidence showing that early life influences have a critical effect on brain connectivity in later childhood and adult life (Batalle et al., 2018). This has key implications for understanding the pathophysiology of neurodevelopmental conditions, such as autism (Hisle-Gorman et al., 2018) or difficulties associated with parturition (Montagna & Nosarti, 2016).

The developing Human Connectome Project (dHCP) was funded by the European Research Council (ERC) and designed to address the need for new and fundamental knowledge about human brain development across the key period from 20 weeks gestation to full term (Edwards et al., 2022). This was achieved through novel and bespoke imaging methods, which were developed and optimized specifically for studying the developing brain with a particular focus on addressing challenges related to subject motion and enabling detailed exploration of brain maturation trajectories. dHCP participants were scanned both *in utero* and after delivery, with some subjects being examined in both states. The imaging data is complemented by rich collateral data including demographic, clinical, and family information to allow new insight into how these factors influence and modulate early brain development. In addition, postnatal saliva samples were obtained for genetic and epigenetic analysis, with participating families invited back when their infants were around 18 months of age for a developmental assessment using standard tests and questionnaires, including eye-tracking studies.

A key priority for the project was that the data be made available to the research community. Neonatal data have been released and described previously (Edwards et al., 2022). Here we describe the fetal data release, providing a summary of the participants, the MRI data acquisition, reconstruction and processing, the collateral data including sociodemographic and neuropsychological outcome data, and the genomic data. We describe available data for each category and how to obtain it. This comprehensive data together with detailed data release notes providing more information, including about the processing pipelines used can be found in Project 3955 on National Institute for Mental Health (NIMH) data repository portal (https://nda.nih.gov/edit_collection.html?id=3955) and general project information is available at http://www.developingconnectome.org/data-release/.

## Methods

### Data Overview

This dataset rests on optimized MR acquisition sequences, novel image reconstruction methods, data processing using state-of-the-art pipelines, packaged up for final public release. The data release contains anatomical MRI (T1 weighted (T1w) and T2 weighted (T2w)), resting state functional MRI (fMRI), diffusion MRI (dMRI), alongside EPI reference scans and field maps for the measured static (B0) and radiofrequency (B1+) magnetic fields. For each modality, the data release includes both the original image data and outputs after the various processing pipelines described below have been applied. For anatomical data, the original multi-slice images are released in addition to motion corrected Slice-to-Volume Reconstructions (SVR), which provide self-consistent isotropic image volumes.

The fetal brain has significantly different tissue properties to the adult brain, including higher water content and incomplete myelination of white matter, resulting in T1 and T2 relaxation times that are generally longer than in the adult brain. Fetal white matter in particular, has significantly longer T1 and T2 times than grey matter (Bhattacharya et al., 2024), and brain anatomy is revealed more clearly on T2 weighted (T2w) images as there is greater contrast compared to T1 weighted (T1w) images (Rutherford, 2009). T2w images are thus treated as the primary data for anatomical segmentation and to provide the anatomical substrates for functional and diffusion analyses. In addition, fetal motion and maternal respiratory motion, dynamic susceptibility changes, and lower signal-to-noise ratio (SNR) in abdominal imaging, introduce specific challenges which are not present in conventional (*ex utero*) brain imaging. To ameliorate the effects of fetal motion during image acquisition, motion-tolerant acquisition and reconstruction approaches were developed as outlined below.

Native reconstructed images from 273 fetuses aged 21 to 38 weeks gestation over 297 sessions are released, including T2w images from 296 sessions, fMRI image series from 275 sessions and dMRI data from 272 sessions. The release also contains 281 sessions with T1w images (50 acquired using a spoiled gradient recalled echo (SPGR) method, 229 acquired using a balanced steady state free precession (bSSFP) method and 2 with data from both methods (*see below*)). Information about the quality control (QC) process for the native data is provided in the notes accompanying the data release.

### Imaging acquisition methods and parameters

All data were acquired on the same Philips Achieva 3T system (R3.2.2 software release) as the dHCP neonatal imaging data. Following a prior study to determine the optimal maternal positioning, subjects were scanned with mothers supine using a 32-channel cardiac coil (Hughes et al., 2021). This position has advantages for coil placement and geometrical stability of the imaging volume and reduced variability between subjects and examinations. Blood pressure and peripheral pulse oximetry were monitored throughout all examinations, which were supervised by midwives. In the event of signs of vasovagal distress, subjects were immediately removed and mothers who were able to continue were repositioned as needed prior to seeking subject consent for re-commencing scanning. Maternal temperature was monitored using an ear thermometer prior to and following MRI.

Fetal MRI at 3T presents many challenges, particularly sensitivity to field inhomogeneity. Consequently, examinations utilised localised image-based static field (B0) shimming (Gaspar et al., 2018), radiofrequency (RF) field (B1+) shimming. For functional, diffusion and T1w-SPGR acquisitions, local RF power scaling was applied to directly optimise the flip angle in the fetal brain while remaining within SAR limits.^1^ Sequences were extensively tested and optimised with (local enhanced) safety constraints relevant to *in utero* scanning; namely, restricting Specific Absorption Rate (SAR) to <50% local torso ∼ <0.8W/kg whole body, peripheral nerve stimulation (PNS) to <60% scanner limit and acoustic noise as measured in an empty bore to <110dBA (Hutter, Price, et al., 2018). Sequences were refined to balance performance with acoustic noise, by determining a bespoke gradient mode for each modality and adapting sequences to avoid mechanical system resonances. The total fetal exam was divided into two sessions each of ∼30 minutes: the first containing collection of multi-band (MB) structural T2w and fMRI data, and the second consisting of structural T1w and MB dMRI data acquisitions. Single-band fMRI and dMRI reference scans, dual-TE B0 and DREAM (Nehrke & Bornert, 2012) B1+ maps were collected in both sessions for calibration purposes. Snapshot, multi-slice (2D) acquisitions were used for all modalities in order to freeze maternal induced motion and spontaneous fetal movements. Parameter values are listed in Table 1.

**Table 1.** Acquisition parameters of the fetal dHCP protocol: core modalities (Price et al., 2019).

|  | **T2w** | **T1w-SPGR** | **T1w-bSSFP** | **fMRI** | **dMRI** |
|---|---|---|---|---|---|
| **Encoding** | ssFSE | SPGR | bSSFP | GE-EPI | SE-GE-EPI |
| **Acq. Res. (mm)** | 1.1x1.1x2.2 (-1.1 gap) | 1.5x1.5x4.0 (-1.2 gap) | 1.5x1.5x4.0 (-1.2 gap) | 2.2x2.2x2.2 | 2.0x2.0x2.0 |
| **Slices** | 48/55 | 42(x2) | 42 | 48 | 54 |
| **In-plane acc.** | Zoom 50% | SENSE = 1.8 | SENSE = 1.8 | SENSE = 1.4 | SENSE = 2.0 |
| **MB factor** | 2 | - | - | 3 | 2 |
| **P.Fourier** | 0.65 | - | - | - | 0.75 |
| **Orientations** | 6 unique | 4 unique | 6 unique | Axial | Axial |
| **TE (ms)** | 250 | 4.6 | 3.6 | 60 | 76/137 |
| **TR (ms)** | 2265+ | 10 | 7.2 | 2200 | 5810 |
| **Shot dur. (ms)** | 1052 | 1230 | 885 | 105.8 | 60.3 |
| **Flip angle (<sup>0</sup>)** | 90/130 | 8 | 35 | 70 | 90/180 |
| **RF shim** | PTx | PTx | PTx | PTx | PTx |
| **Power scaling** | Global | Local | Global | Local | Local |
| **B0 shim** | IB brain | Default | IB brain | IB brain | IB brain |
| **Volumes** | 6 | 8 | 8 | 350 | 141 |
| **Scan times (min)** | ~12 | ~8 | ~8 | 13.0 | 13.75 |
| **Preparation pulses** | SPIR, Tip-back pulse | IR (TI=2.72s) | IR (TI=2.73s) | SPIR, OVS band | SPIR, grad. reversal |

**Table 2.**
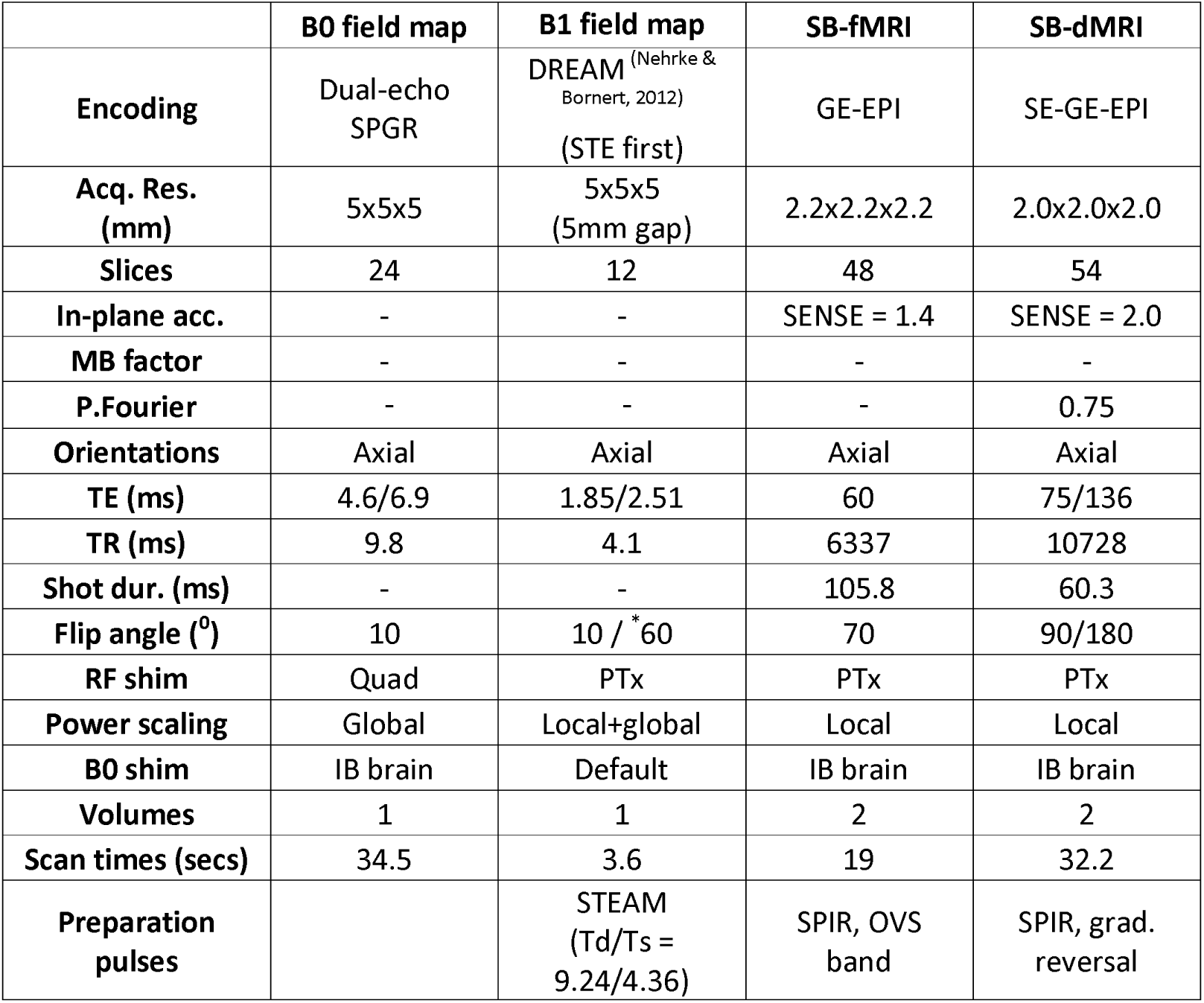
Acquisition parameters of the fetal dHCP protocol: calibration scans in data release.

**Structural T2w data** was acquired from 6 stacks, each with a different orientation, centred to the fetal brain using a zoomed multi-band 2 (MB2) single-shot Fast Spin Echo (ssFSE) sequence, with a MB tip-back preparation pulse for increased SNR efficiency (Price et al., 2018). Localised B1+ shimming was utilised without local power scaling due to SAR constraints on minimum TR. B0 shimming was necessary due to the field sensitivity of the tip-back preparation.

**Structural T1w data** was collected from 8 stacks at 6 unique views (2 views collected twice) using an inversion-recovery acquisition with preparation that inverted an individual double slice-width slab for each slice to provide optimised contrast with increased robustness to movement between inversion and readout. Initially data was collected using a SPGR readout (Ferrazzi et al., 2018), but following further testing this was replaced with bSSFP readout to improve SNR (Nehrke & Bornert, 2012). In both cases the scanner software was modified to allow efficient temporal interleaving of the inversion pulses and readout echo trains (Ferrazzi et al., 2018). B1+ shimming was utilised with and without local power scaling for T1w-SPGR and T1w-bSSFP, respectively (Price et al., 2019).

**Functional data** collecting 350 volumes utilised MB3 slice acceleration and echo planar imaging (EPI) readout with SENSE reconstruction (Karolis et al., 2025). Only moderate in-plane acceleration (x1.4) was used, since a longer echo time benefits the MB-unfolding, due to fast decay of maternal tissues in the extended field-of-view, and potentially enhances BOLD contrast (Ferrazzi, 2016).

**Diffusion data** consisting of 141 volumes with 2 shells (15 b=0 s/mm, 46 b=400 s/mm, and 80 b=1000 s/mm) was acquired using a spin- and field-echo (SAFE) sequence for dynamic distortion correction and data collection efficiency (Christiaens et al., 2019; Cordero-Grande et al., 2018). The choice of b-values and number of directions per shell were determined after empirical testing using the methods described in Tournier et al (Tournier et al., 2020) with the shell values finally selected to match the lower shells of the dHCP neonatal data acquisition (Edwards et al., 2022). As for the neonatal acquisition, the sensitisation directions were set using an electrostatic repulsion model, and acquisition order was interleaved between the shells to even out thermal demands on the gradient system (Hutter, Tournier, et al., 2018).

Both functional and diffusion data acquisitions deployed B1+ and B0 shimming and local power scaling.

### Reconstruction and processing pipelines

Reconstruction and processing pipelines for all three MRI modalities (anatomical, diffusion and functional imaging) have been developed specifically for the dHCP fetal data. The outputs of these pipelines together with links to open-source code for different elements of the pipelines are supplied as part of the data release. Details of the individual pipelines are given below.

### Structural T2w reconstruction pipeline

Original images from motion-corrupted ssFSE stacks were reconstructed with multi-band SENSE (MB-SENSE). Then three-dimensional (3D) T2w brain images were reconstructed from these to 1.0mm isotropic resolution using a dedicated fully automated SVR pipeline (Cordero-grande, Price, et al., 2019). The pipeline includes denoising, correction of inhomogeneity using B1+ calibrations, V-net localisation of the fetal brain, followed by several interleaved iterations of SVR and super resolution reconstruction, and is implemented in Matlab (The Mathworks Inc, Natick MA, USA). The 3D reconstructed images were reoriented to the standard radiological planes using a dedicated transformer network for fetal brain MRI (Wright et al., 2018). This was followed by cubic spline interpolation to 0.5×0.5×0.5mm resolution, which is more convenient for further image analysis steps.

### Structural T1w reconstruction pipeline

The 3D T1w brain images were reconstructed to 0.8mm resolution using an in-house fully automated pipeline based on the classical 3D SVR reconstruction method (Kuklisova-Murgasova et al., 2012) in SVRTK (https://github.com/SVRTK/SVRTK) with additional 3D UNet brain masking and landmark-based reorientation to the standard radiological space in auto-SVRTK (https://github.com/SVRTK/auto-proc-svrtk).

### fMRI reconstruction pipeline

The data underwent MB-SENSE image reconstruction, dynamic shot-by-shot B0 field correction by iterative least-squares phase unwrapping (Cordero-Grande et al., 2018) (see source code at https://github.com/mriphysics/fetalPhaseEPI/releases/tag/1.0.0) and SVR motion correction. Following spatial corrections, a temporal denoising model optimised for fetal data signal recovery was applied (Karolis et al., 2025; Karolis et al., 2023).

### dMRI reconstruction pipeline

The 3D dMRI fetal brain images were reconstructed with MB-SENSE to 2.0mm resolution using an in-house pipeline implemented in Matlab and the MRtrix3 toolbox (Tournier et al., 2019). It includes Generalized Singular Value Shrinkage (GSVS) MP-PCA image denoising (Cordero-Grande, Christiaens, et al., 2019) (source code: https://github.com/mriphysics/complexSVDShrinkageDWI), phase-based data dynamic distortion correction of susceptibility-induced B0 field changes based on iterative least-squares (Zhao et al., 2020) leveraging dual echo data for removing the underlying unpredictable phase due to motion during the diffusion sensitizing gradients (Cordero-Grande et al., 2018), and SVR motion correction of the dual-echo data based on a multi-shell spherical harmonics and radial decomposition (SHARD) representation (Christiaens, Cordero-Grande, et al., 2021; Christiaens, Pietsch, et al., 2021).

### Volumetric alignment between the modalities

The T1w and dMRI images were semi-automatically co-aligned to T2w images using in-house deep learning landmark-based global rigid alignment followed by boundary-based registration implemented in FSL FLIRT with local absolute differences as a cost function (Smith et al., 2004). Alignment of fMRI to T2w was performed using rigid whole-brain FSL FLIRT linear registration (using global search and mutual information as a cost function), refined using by boundary-based registration, using the same cost function as for the dMRI data. Alignment from the subject T2-weighted to the dHCP atlas space (see below) was performed using affine FSL FLIRT, followed by nonlinear multi-channel registration guided by T2w images and cortical probability using ANTs (Tustison et al., 2021). The output transformations include affine and nonlinear warp components (Karolis et al., 2023).

### Anatomical segmentation and surface generation pipelines

The T2w SVR images were segmented into 43 main tissue classes with left/right hemisphere separation using the deep learning Multi-BOUNTI pipeline (Uus et al., 2026) (https://github.com/SVRTK/perinatal-brain-mri-analysis). All parcellation outputs underwent a QC process by a neuroimaging expert (VK) (See the notes accompanying the data release for details).

Surface generation was performed using a modified, fetal anatomy-adapted CoTAN network (Ma et al., 2023), referred to here as FetalSeg-CoTAN (https://github.com/irinagrigorescu/FetalSeg-CoTAN). The network utilizes a template-based deformation framework, in which subject-specific cortical surfaces are generated by deforming an initial template mesh, defined as a sixth-order regular icosphere containing 40,962 vertices and 81,920 faces in a 36-week fetal atlas space. To optimize computational efficiency while preserving key anatomical boundaries, the input to FetalSeg-CoTAN consists of simplified multi-BOUNTI segmentations, where the original 43 anatomical labels are consolidated into 5 tissue classes (white matter, cortical grey matter, cerebrospinal fluid, brainstem, and cerebellum). Operating on these simplified inputs, the network predicts subject-specific vertex displacements via learned, gestational age-conditioned deformations.

Final outputs are provided in subject (native) space, consisting of separate surfaces for each hemisphere (left and right white matter and pial surfaces). In addition to the primary surfaces, we generate mid-thickness, inflated, and very inflated surfaces, as well as spherical representations using FreeSurfer v8.1.0 (Fischl, 2012). To facilitate downstream analysis, we also compute morphometric maps of cortical thickness, sulcal depth and mean curvature, as well as cortical parcellations. All outputs underwent a QC process (AB, NC, MD, MD, IG, HS, WW, CZ) (See the notes accompanying the data release for details). In summary, 283 (96%) reconstructed surfaces passed QC and were scored good or excellent.

Technical note: As one of the preliminary results of data processing, the initial data release also included outputs of the optimised Draw-EM segmentation pipeline previously used in the neonatal dHCP release (Makropoulos et al., 2018) (https://github.com/DevelopingHCP/structural-pipeline/tree/feature/fetal_pipeline). However, the Draw-EM labels and surfaces have suboptimal quality and are not recommended for detailed analysis.

### Spatio-temporal Fetal Brain MRI Atlases

Similarly to the dHCP neonatal structural atlases (Schuh et al., 2018; Schuh et al., 2014) (Uus, Kyriakopoulou, et al., 2023) (Bozek, Makropoulos, Schuh, Fitzgibbon, Wright, & Glasser, 2018) the fetal dHCP datasets were used to create volumetric and surface spatio-temporal MRI atlases of fetal brain development (Uus, Hall, et al., 2023).

For the volumetric atlas, in addition to structural T2w and T1w data, the following dMRI measures (extracted in MRtrix3) were included: fractional anisotropy (FA), mean diffusivity (MD), radial diffusivity (RD), orientation distribution functions (ODF) and average b=1000 s/mm shell (Figure 6). Prior to atlas construction, all modalities were aligned to individual subject bias-field corrected T2w space. The selection criteria for data to be used for atlas construction were: no reported congenital or acquired brain anomalies, image quality and registration performance QC based on visual assessment. All images were visually checked for the presence of artefacts and sufficient visibility of all brain structures. The atlas was created using the unbiased atlas generation tool (Schuh et al., 2018) as a baseline with diffeomorphic registration using the MIRTK toolbox that was previously used for the structural neonatal dHCP atlas (Uus, Kyriakopoulou, et al., 2023). This process consisted of the following steps: T2w+cortex label guided nonlinear registration, local normalised cross-correlation similarity metric, one affine and 3 nonlinear iterations, 1 week temporal kernel followed by additional manual refinement of the registration step for individual subjects with case-specific adapted settings in order to improve registration performance for the cortical ROI. The resulting transformations and weighted averaging (with Laplacian sharpening) were used to produce averaged images with 0.5mm isotropic resolution in the reference space for all channels. The output atlas has 16 timepoints from 21 to 36 weeks GA for all 7 channels.

The fetal surface atlas was generated using the previously described framework for the dHCP neonatal surface template generation (Bozek, Makropoulos, Schuh, Fitzgibbon, Wright, Glasser, et al., 2018; Liang et al., 2025; Robinson et al., 2018; Serag et al., 2012; Williams et al., 2023). All registrations during template generation were performed using Multimodal Surface Matching (MSM). Individual fetal surfaces were first rigidly aligned to the HCP template space using sulcal depth to establish coarse folding correspondence. Week-specific templates were then generated using adaptive kernel weighting to account for the uneven distribution of gestational ages while preserving consistent anatomical detail across weekly templates. Because sulcal depth mainly captures large-scale folding patterns, the templates were subsequently refined using curvature to improve alignment of finer gyral and sulcal features. Registration drift introduced during iterative refinement was corrected before generating the final week-specific surface and metric templates. Left–right vertex correspondence was then achieved using MSM registration, enabling direct comparison of homologous cortical locations between hemispheres. Finally, consecutive weekly templates were registered and the resulting deformations concatenated to map all fetal surfaces into a common 36-week template space.

### Governance and Access

The study was approved by the Riverside NHS Research Ethics Committee (reference number: 14/LO/1169) and written parental consent was obtained in every case for imaging, collection of other data and open data release of the de-identified pseudo-anonymized data. The main imaging data, essential metadata and collateral data are available after accepting a data sharing agreement. Downloaded data should not be passed on to third parties outside the original accessing research group, and no attempt should be made to de-anonymize the data.

The primary long-term site for curation and access of the full data release is the National Institute for Mental Health (NIMH) data repository portal at https://nda.nih.gov/edit_collection.html?id=3955. Details on data access, including detailed information about data organisation, file naming conventions and links to pipeline source code, are available on the portal. The atlases are publicly available online at the dHCP data repository: https://gin.g-node.org/kcl_cdb/fetal_brain_mri_atlashttps://gin.g-node.org/kcl_cdb/.

## Results

### Participants

Mothers were recruited with the following inclusion and exclusion criteria. Pregnant women were eligible for inclusion at 20–42 weeks’ gestation (estimated from the last menstrual period and confirmed where possible by early ultrasound). The study included pregnancies with normally developing fetuses as well as those at increased risk of neurodevelopmental disorder, including genetic risk and potential environmental factors such as fetal growth restriction. Exclusion criteria included contraindications to MRI, including incompatible implanted metallic devices, and language difficulties preventing adequate communication and informed consent. Participants could withdraw if an exclusion criterion developed during the study or if they chose to withdraw. The distributions of gestational age at scan (range 20+6 - 38+2 weeks) are shown in figure 1.

**Figure 1.**
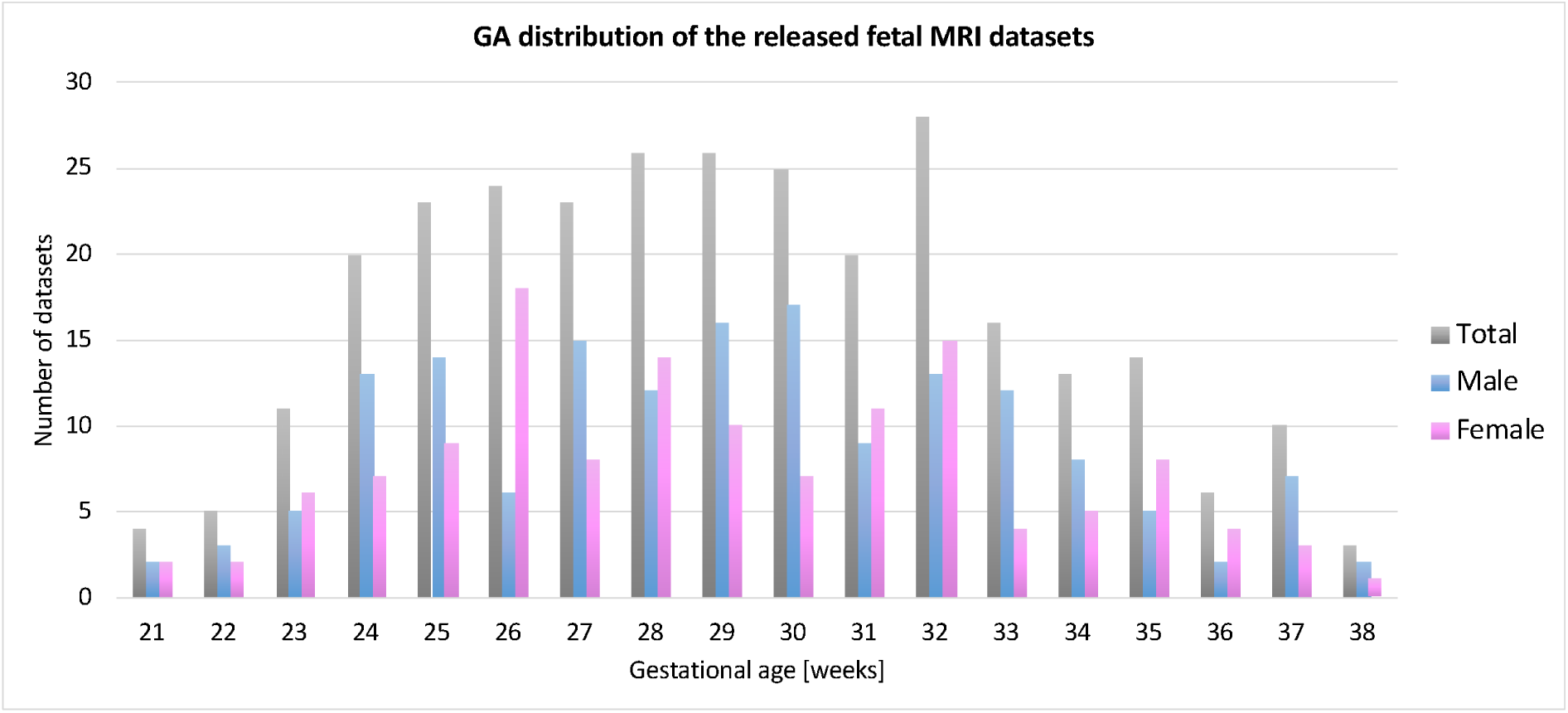
Histograms showing gestational age distribution of fetuses with released MR datasets.

The study population includes 273 subjects with one subject not having a successful T2w SVR reconstruction (the native T2w images are included and this subject had usable data in all other modalities). All the anatomical images were reviewed by an expert perinatal neuroradiologist (MAR), and radiologic scores are included in the released data.

### Exemplar imaging datasets

Figures 2 – 6 show exemplars from individual subjects to provide an indication of the released imaging data.

**Figure 2.**
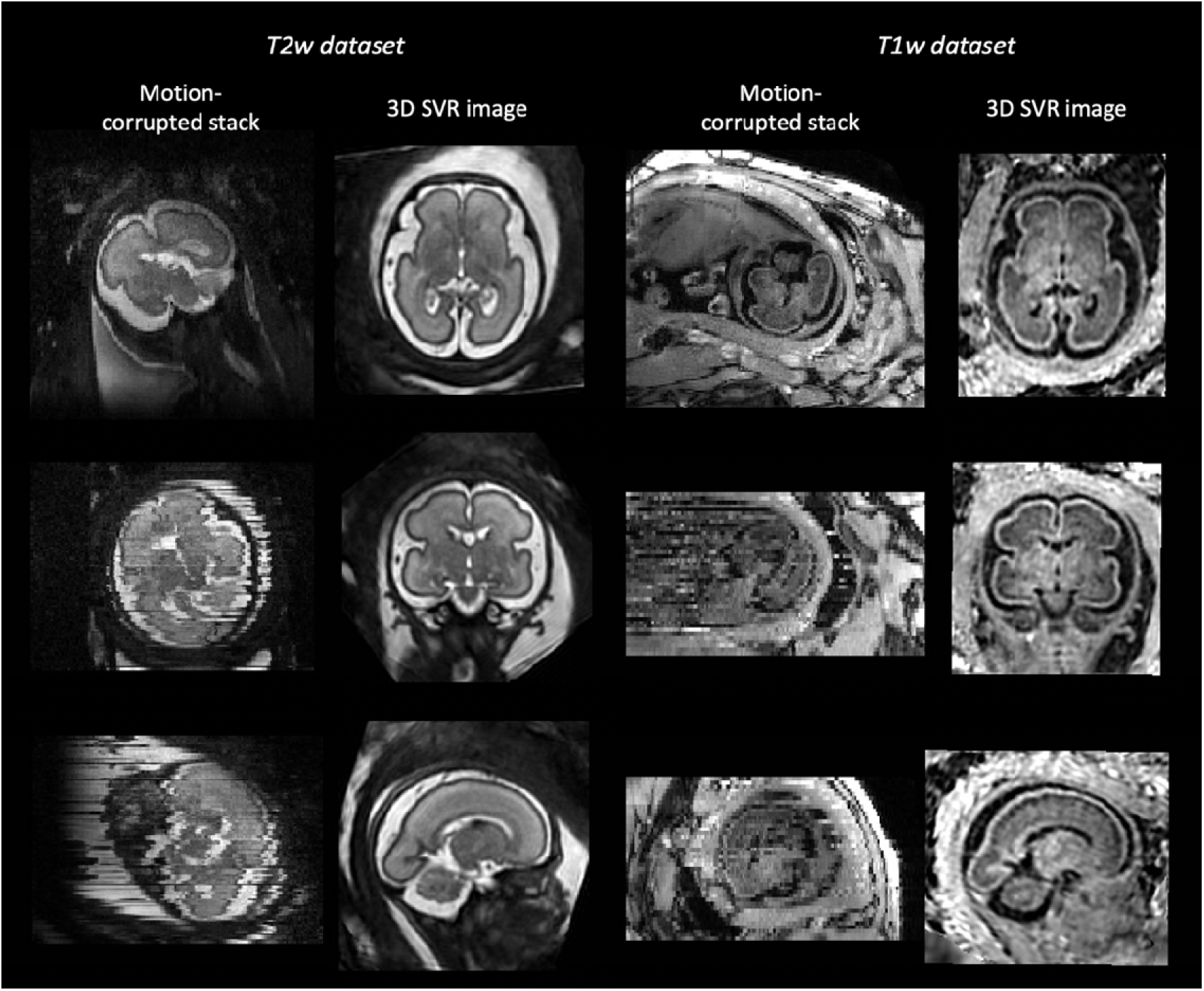
Anatomical T2w (left) and T1w (right) images for one participant (GA = 26w4d). Rows show orthogonal views (Transverse, Coronal and Sagittal). For each contrast, one input motion-corrupted stack is shown on the left and the output motion-corrected 3D SVR reconstructed images are shown on the right.

**Figure 3.**
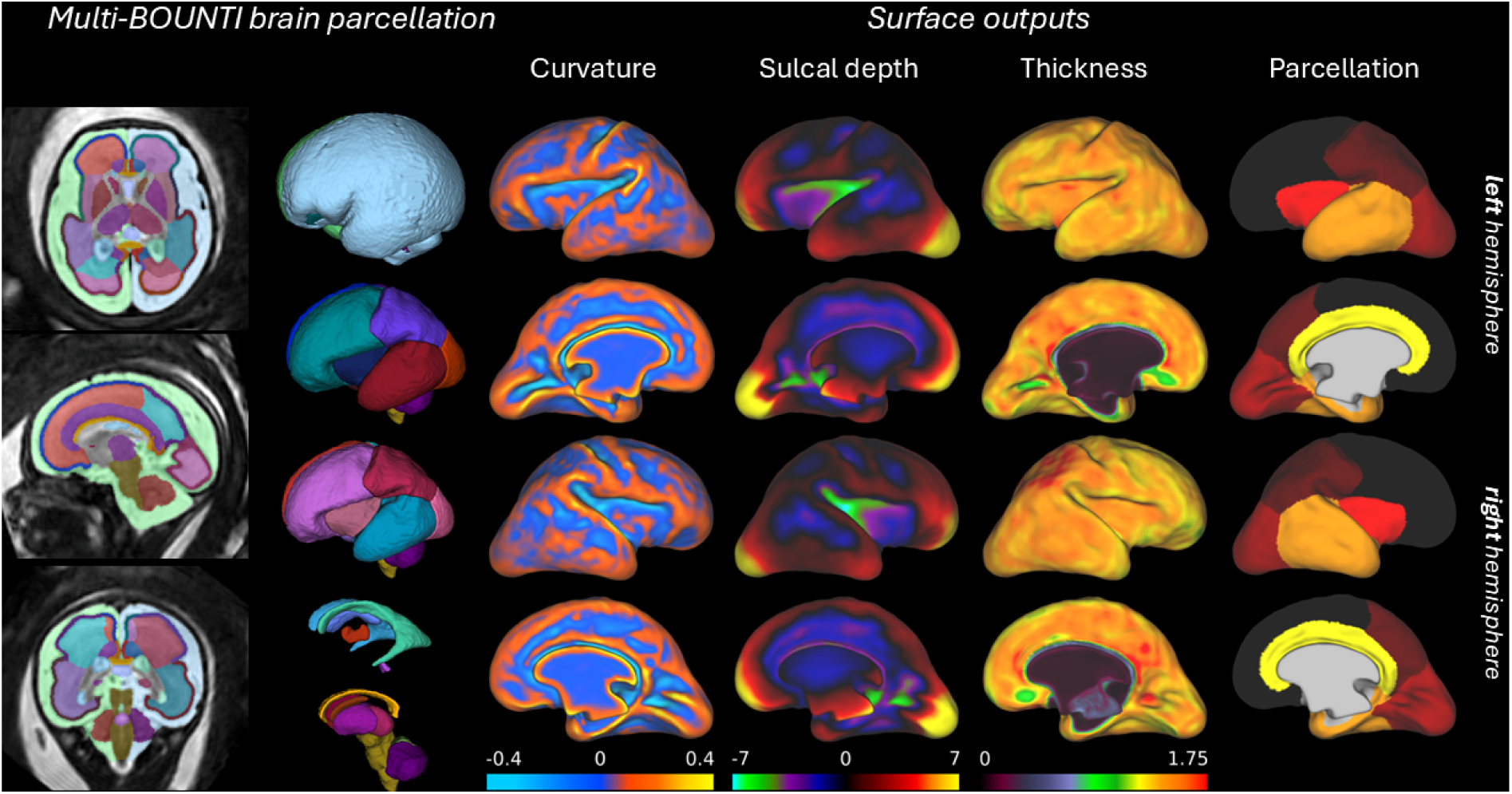
Multi-BOUNTI Tissue segmentation and surface maps for the same subject as figure 2 (GA = 26w4d) *-* generated *u*sing the automated deep learning Multi-BOUNTI pipeline. Anatomical images were segmented into 43tissue classes. The surface reconstruction pipeline outputs include curvature, sulcal depth, cortical thickness, and cortex parcellation.

**Figure 4.**
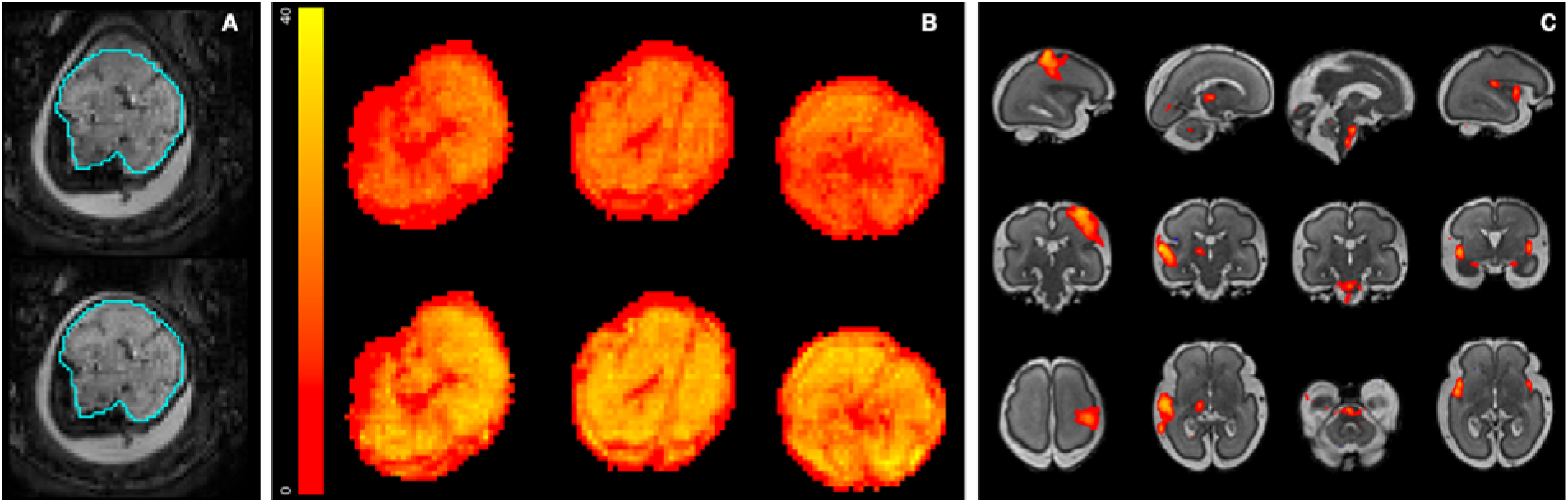
Resting state fMRI data for the same participant as in figure 2 (GA = 26w4d). A) Single volume from the fMRI time series acquisition showing the effect of dynamic distortion corrections; top – before distortion correction, bottom – after distortion correction. B) Temporal SNR before (top row) and after (bottom row) temporal denoising. C) Example resting state networks of probable biological origin.

**Figure 5.**
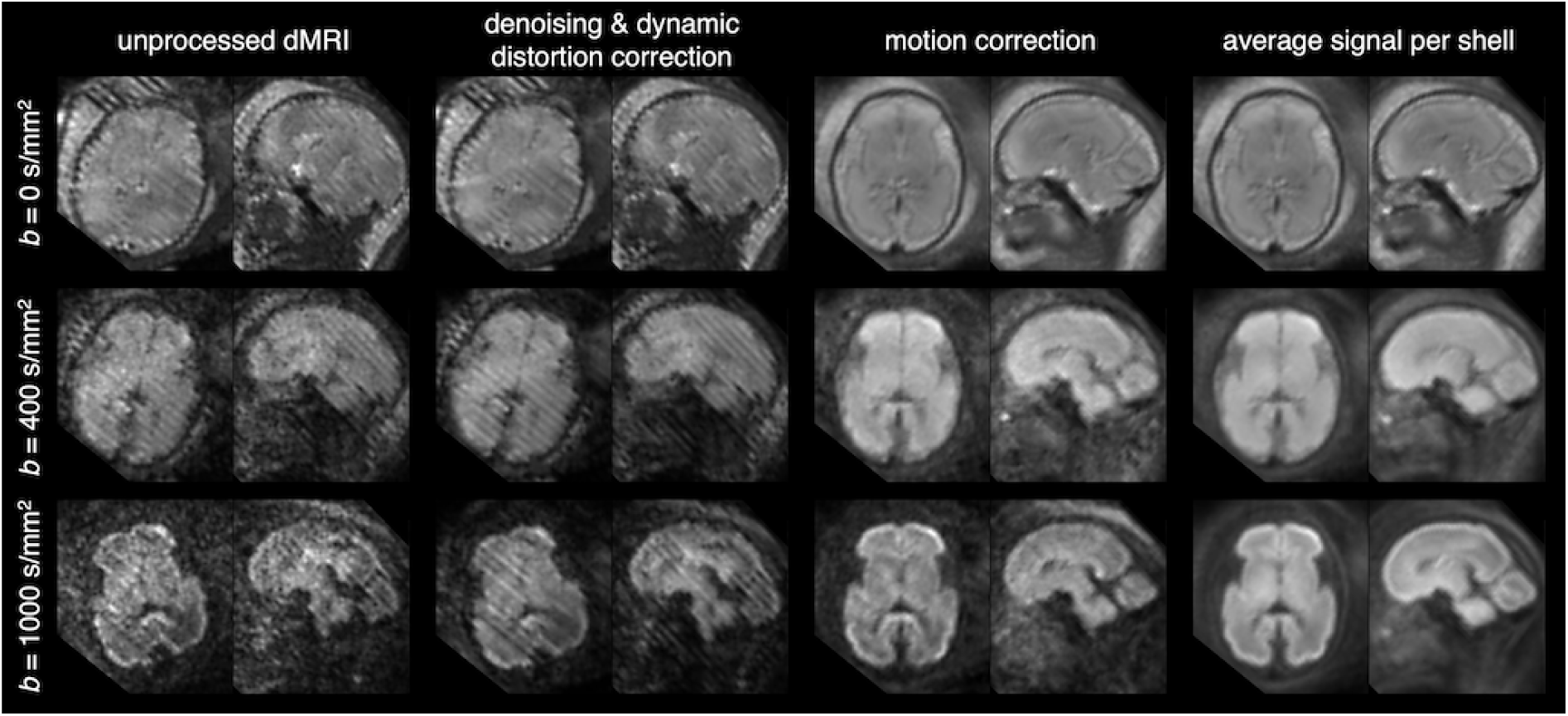
dMRI data for one participant (GA = 28w2d): Transverse and sagittal views for each b-value (rows). Left column: typical slice stacks. Second column: corresponding slice stacks after slice-wise denoising and distortion correction. Third column: predicted signal in corresponding volumes after motion correction with SHARD-SVR. Right column: mean signal across all diffusion encoding directions in each shell (each image normalized separately for visualisation).

**Figure 6.**
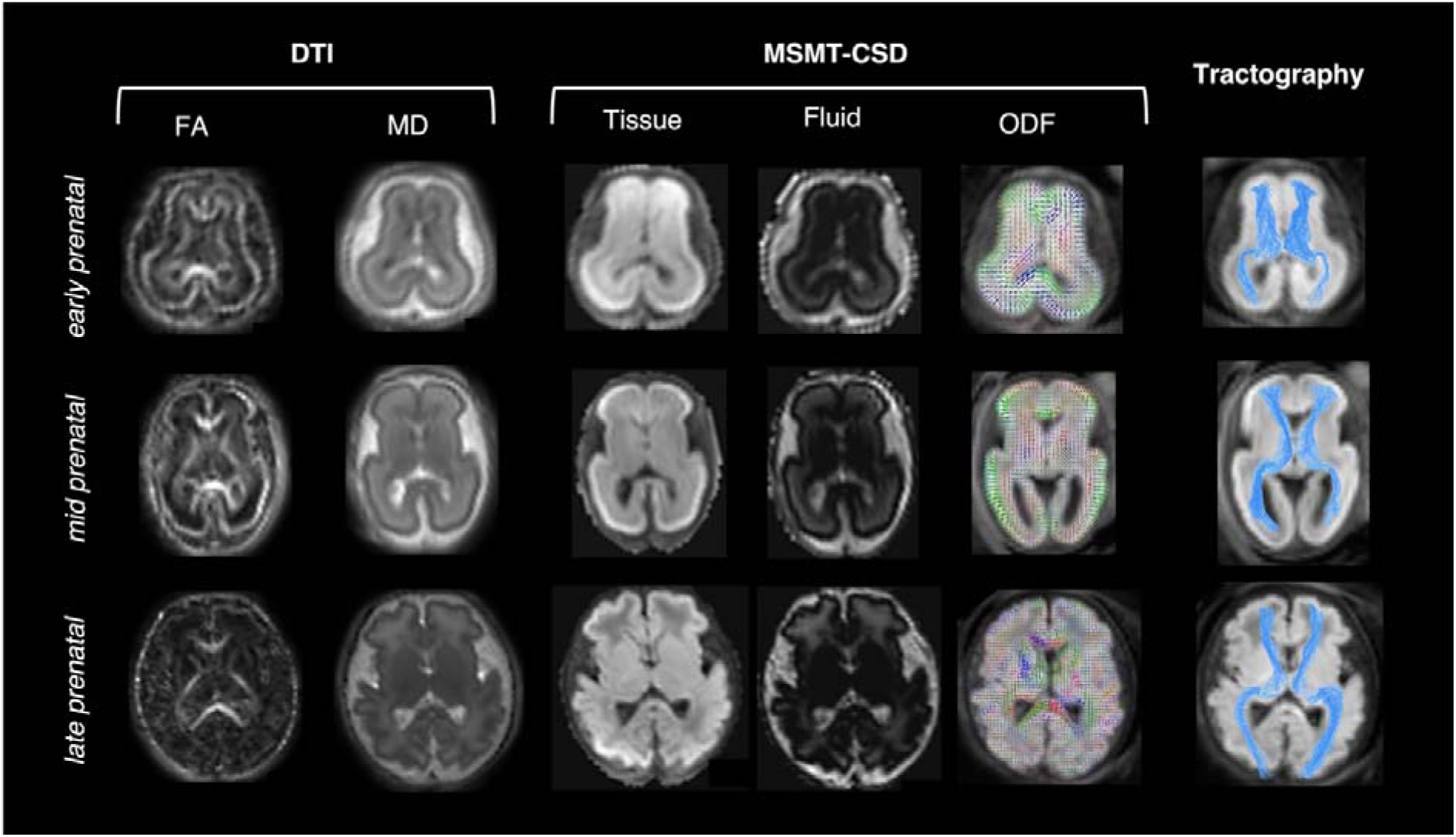
dMRI metrics extracted for 3 participants at early (22 weeks), Mid (28 weeks) and late (33 weeks) gestational age. Left: Fractional Anisotropy (FA) and Mean Diffusivity (MD) calculated using a diffusion tensor model; Centre: tissue, fluid and Orientation Distribution Function (ODF) maps; Right: Tractography (Wilson et al., 2021) calculated from a Multi-Shell Multi-Tissue Constrained Spherical Decomposition (MSMT-CSD) model (Jeurissen et al., 2014; Pietsch et al., 2019).

### Template reference space

Figures 7-9 show the two available dHCP atlases. The volumetric atlas for the whole GA range is summarised in one-week intervals in figure 7. The contributing data was from a total of 187 datasets using 187 T2w, 133 T1w, 135 DWI scans from 21-37 weeks GA. The constant temporal kernel with standard deviation of 1 week ensured spatial continuity between timepoints. Figure 8 shows a single week view in more detail. This atlas (Uus, Kyriakopoulou, et al., 2023) is available at https://gin.g-node.org/kcl_cdb/fetal_brain_mri_atlas.

**Figure 7:**
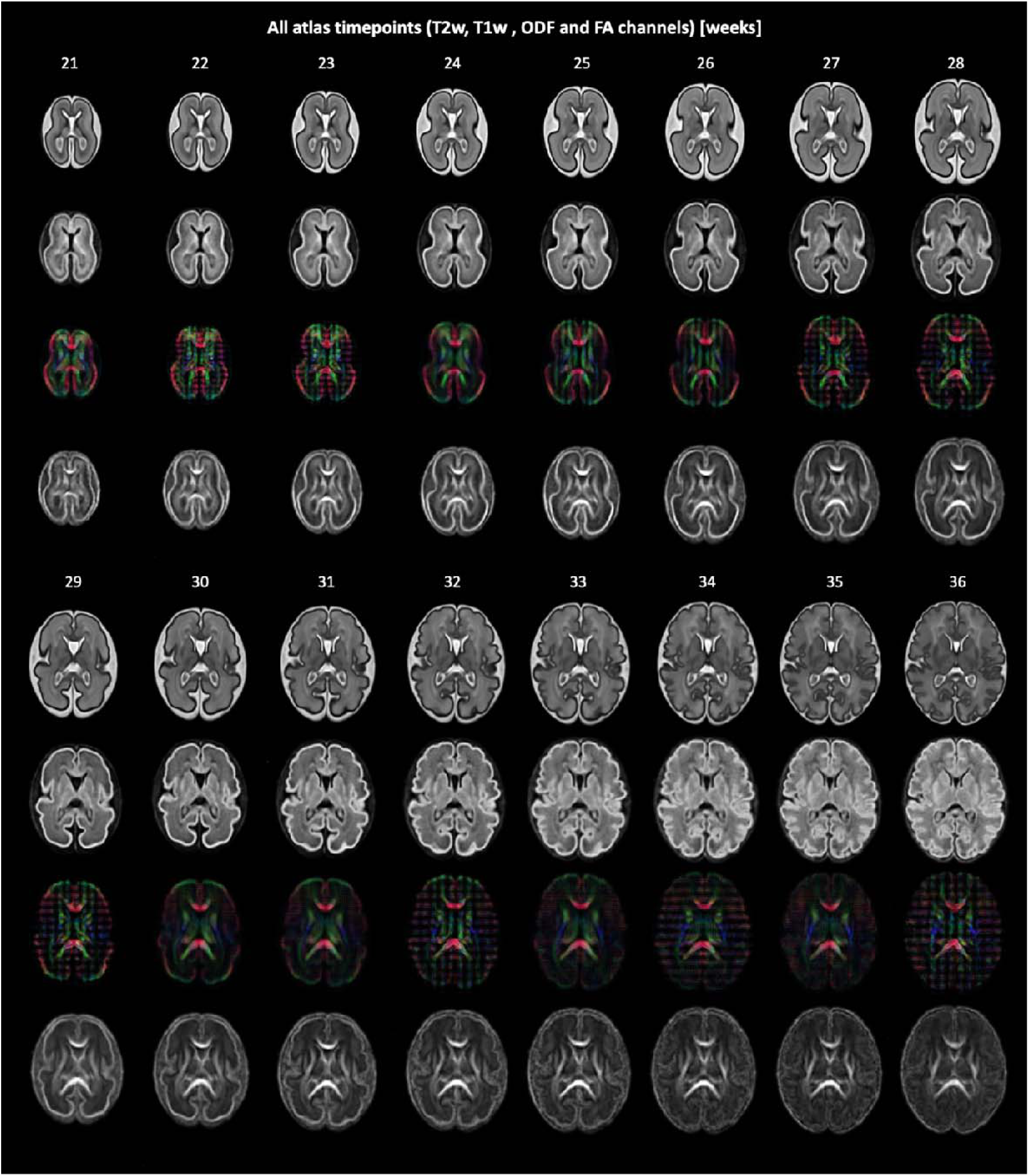
T2w, T1w, ODF and FA channels of the atlas at 21-36 weeks GA timepoints.

**Figure 8:**
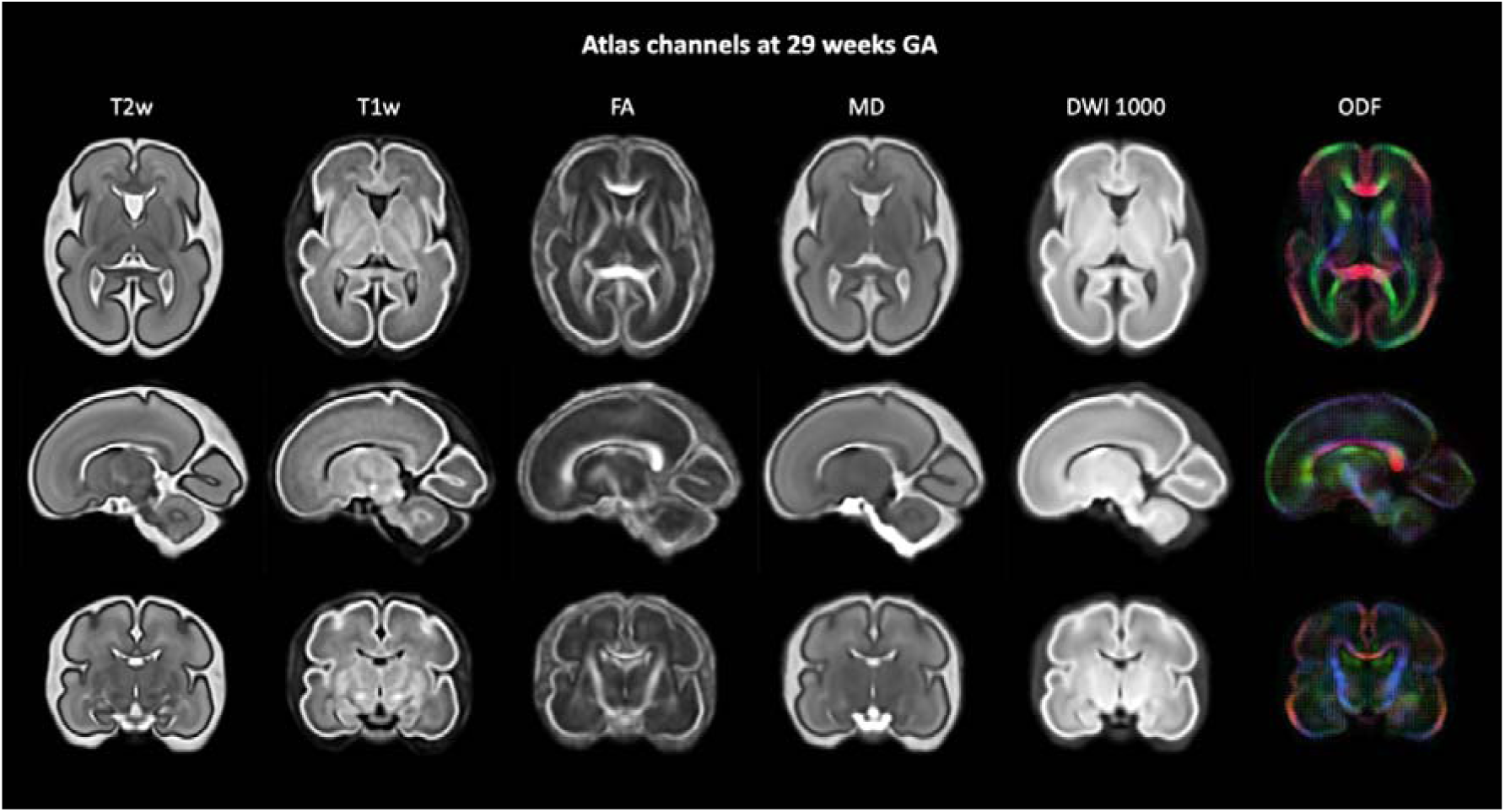
Transverse, Sagittal and Coronal views for the atlas at 29 weeks GA.

**Figure 9.**
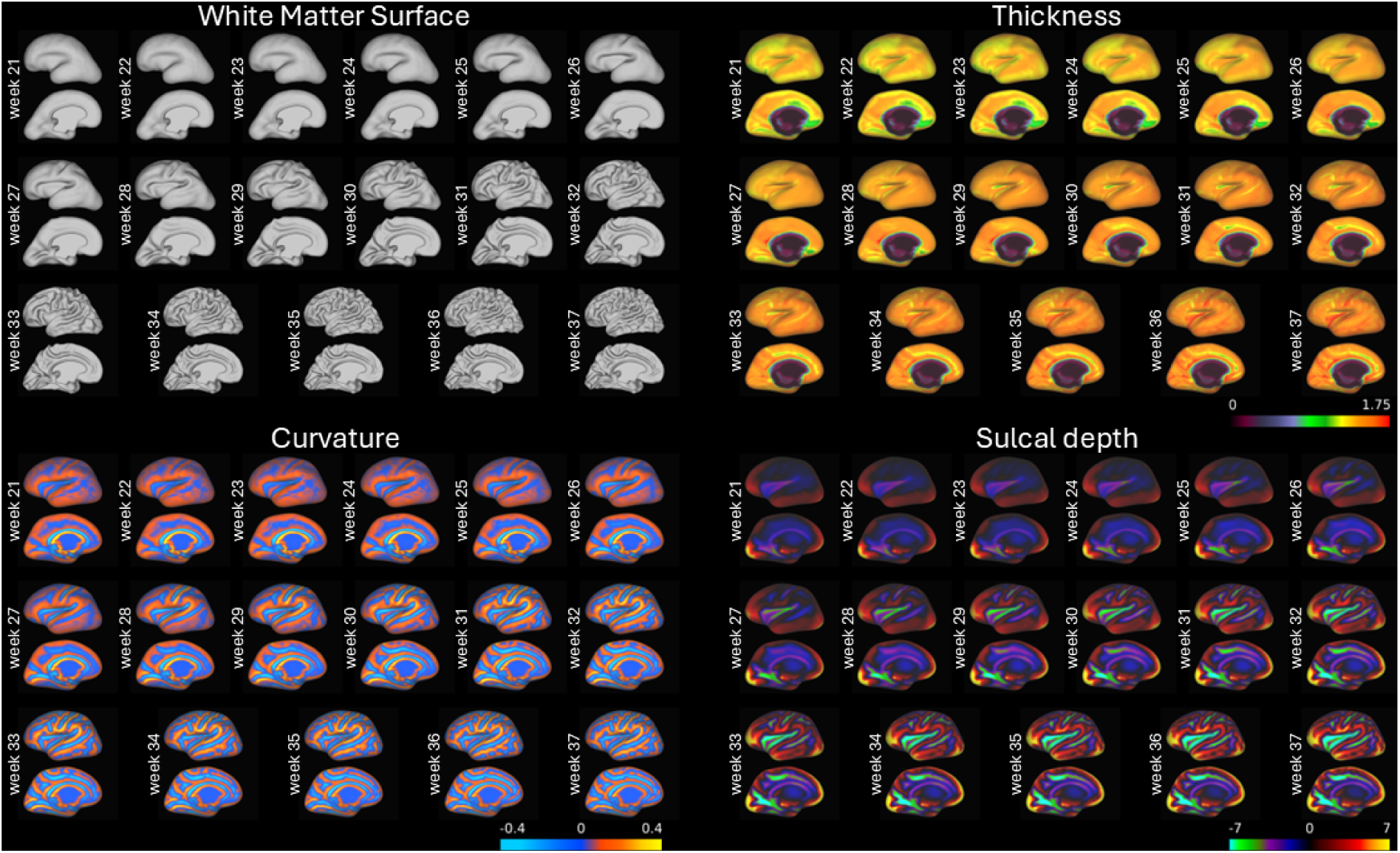
Spatio-temporal dHCP surface atlas summarising 4 channels of information across gestational age.

**Figure 10:**
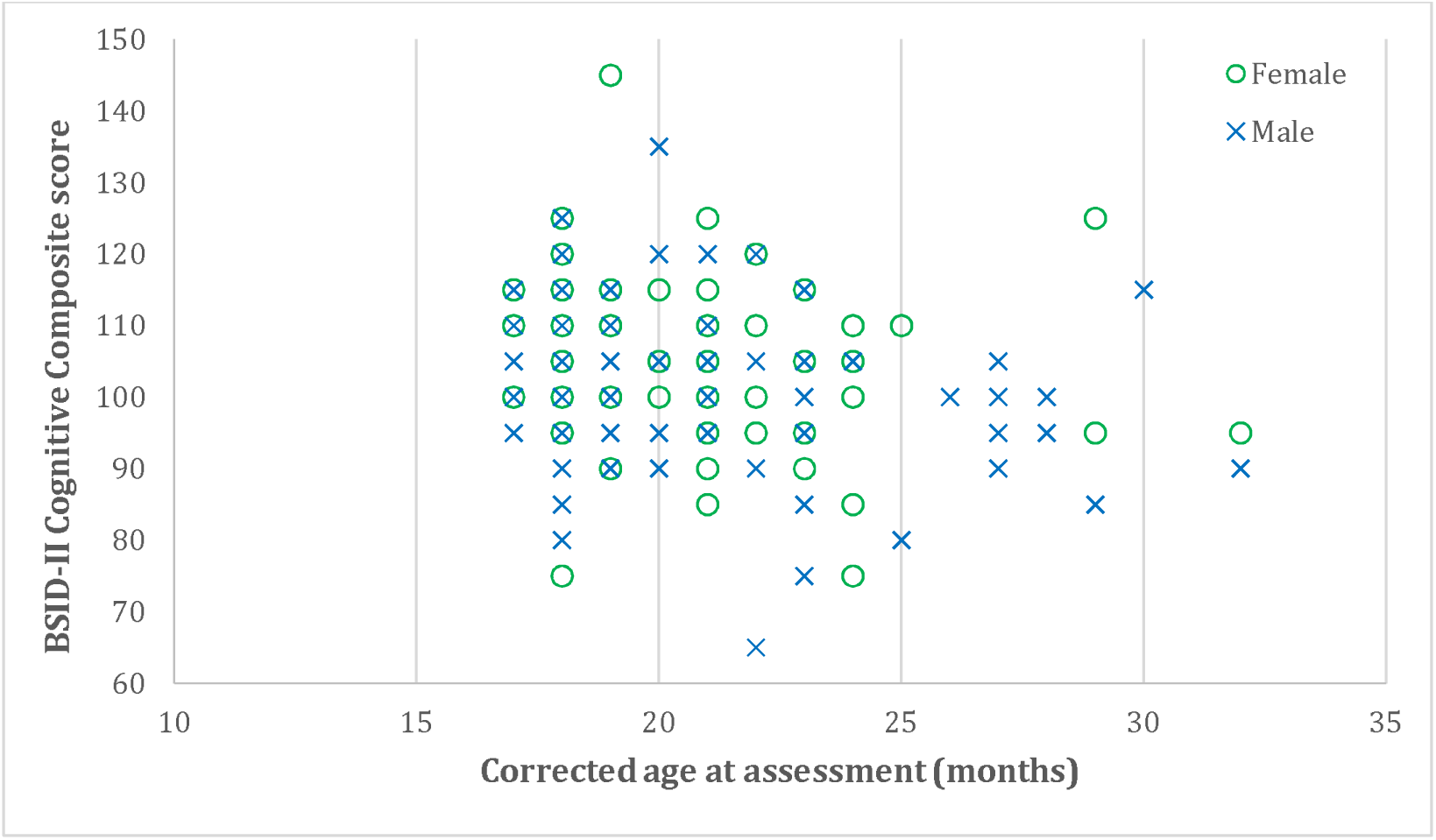
Scatterplot Showing age of assessment and Bayley III cognitive scores for boys and girls (Albers & Grieve, 2007).

Figure 9 shows the spatio-temporal surface template, the first atlas of this type created using a comprehensive dataset. Apart from surface meshes, the atlas includes measures of cortical thickness, sulcul depths and curvature. The atlas is available at: https://gin.g-node.org/kcl_cdb/dhcp_fetal_brain_surface_atlas.

#### Collateral Data

A broad spread of demographic and other data is available, although practical constraints including the COVID-19 pandemic led to a certain amount of missing data. A data codebook can be accessed in the NIMH database (https://nda.nih.gov/edit_collection.html?id=3955) and provides a listing with descriptions of the variables. The data sets include the following categories of data:

### Demographic, Family and Clinical data

#### Demographic Data for Parents

age at conception; ethnicity according to UK census categories; highest age enrolled in full-time education; occupation. This data is collected at enrolment and again at the infant 18-month neurodevelopmental assessment.

#### Mother’s past medical history

height, weight, body mass index (BMI); blood group; history of medical conditions prior to the pregnancy; smoking, alcohol, and recreational drug use; injury during the pregnancy.

#### Mother’s obstetric history

previous pregnancies; number of live births; number of miscarriages; previous premature birth; current pregnancy type, mode of conception (natural or IVF); pregnancy number; late pregnancy and labour/delivery history for the pregnancy.

#### Mental health history

self-reported by mother at enrolment and self-reported by both parents at the 18-month assessment, including any history of parental psychiatric problems and how treated; parental history of attention deficit hyperactivity disorder (ADHD), bipolar disease, autistic spectrum disorder (ASD) or schizophrenia; ASD or ADHD in proband’s siblings; close relatives with history of ASD, ADHD, bipolar disease, or schizophrenia.

### Neurodevelopmental and Neurocognitive testing at 18 months

A series of standardized age-appropriate child-centered assessments, parent-report questionnaires, and gaze-tracking tasks were used to provide a targeted overview of toddlers’ development. These measures were chosen to capture individual differences along a typical-to- atypical continuum, to probe associations between early imaging features and emerging behavioral outcomes and to provide normative reference data for future studies.

A total of 185 children (53% male; n = 98: 47% female; n = 87) attended for follow-up assessment, planned for 18 months corrected age, but affected by the COVID-19 pandemic, so that median (range) of assessment was 19 months + 8 days (range 17 + 18–32 + 22). Parents of a further 16 infants (75%; male n = 12; 25% female; n = 4) completed parent-report questionnaires only. Completion rates for broad components of this assessment are shown in Table 3.

**Table 3.** Completion rates for later infant neurodevelopmental assessments and questionnaires for the 273 recruited fetuses.

| **Neurodevelopmental assessment/Questionnaire (n = 273)** | **Number (%)** |
|---|---|
| Bayley III: cognitive, language, motor neurodevelopmental variables (Albers & Grieve, 2007). | 185 (68%) |
| Hammersmith Infant Neurological Examination: total score (Haataja et al., 1999) | 185 (68%) |
| Early Childhood Behavioral Questionnaire (ECBQ)(Putnam et al., 2006) | 196 (72%) |
| Child Behavioral Checklist (CBCL) (Achenbach & Rescorla, 2000) | 197 (72%) |
| Quantitative Checklist for Autism in Toddlers (Q-CHAT)(Putnam et al., 2006) | 195 (71%) |
| Cognitively Stimulating Parenting Scale (CSPS)(Allison, Baron-Cohen, Wheelwright, Charman, Richler, Pasco, et al., 2008) | 195 (71%) |
| Parenting Scale: primary caregivers' laxness, over reactivity, verbosity(D.S. Arnold et al., 1993) | 196 (72%) |
| Parenting Scale: secondary caregivers' laxness, over reactivity, verbosity(D.S. Arnold et al., 1993) | 179 (66%) |

#### Neurological Examination of Infant/Child

26 non-age dependent items were used to assess cranial nerve function, posture, movements, tone and reflexes at 18 months. (Haataja et al., 1999).

#### Bayley III cognitive, language, motor neurodevelopmental variables

Toddlers’ cognitive, language (receptive and expressive), and motor abilities (gross and fine) were assessed using the Bayley Scales of Infant and Toddler Development, Third Edition (Bayley-III). Composite scores for each domain are age-normed standardized scores with a mean of 100 and standard deviation of 15, ranging from 40 to 160.

#### Early Childhood Behaviour Questionnaire (ECBQ)

this measures dimensions of temperament, referring to individual differences in reactivity and self-regulation (Putnam & Rothbart, 2006). The ECBQ describes three broad scales: Surgency, characterized by impulsivity, intense pleasure seeking and high activity levels; Negative Affectivity, which refers to the disposition to experience aversive affective states, such as anger, fear, anxiety, shame and disgust; and Effortful Control, which refers to the capacity to inhibit/activate a behavioural response by focusing attention.

#### Child Behavioral Checklist for Ages 1.5 to 5 (CBCL)

which is a 100-item measure on the frequency of behavioral and emotional problems in young children (Achenbach & Ruffle, 2000). The CBCL yields scores for seven problem behavior syndrome subscales: Emotionally Reactive, Anxious/ Depressed, Somatic Complaints, Withdrawn, Sleep Problems, Attention Problems, and Aggressive Behaviour. Scores are also derived for Externalizing Problems, Internalizing Problems and Total Problems.

#### Quantitative Checklist for Autism in Toddlers (Q-CHAT)

a 25-item questionnaire designed to assess potential autistic traits in children (Allison, Baron-Cohen, Wheelwright, Charman, Richler, & Pasco, 2008).

#### Cognitively Stimulating Parenting Scale (CSPS)

adapted from (Wolke et al., 2013), which assesses the availability and variety of experiences that promote cognitive stimulation in the home. This includes availability of educational toys, parental interactions such as teaching words or reading stories, and cognitively stimulating activities such as family excursions. The version of the CSPS used here was updated to include four items now widely used by toddlers (i.e., iPhone and Apps) (Bonthrone et al., 2021). Scores from the 28 items included in the CSPS can be aggregated to provide an overall cognitively stimulating parenting score.

#### Parenting Scale

a 30-item rating scale that measures dysfunctional parenting in discipline situations (D. S. Arnold et al., 1993).(D. S. Arnold et al., 1993). Parents are asked to indicate their tendency to use specific discipline practices using a 7-point scale. The Parenting Scale identifies three different suboptimal parenting styles, as well as a total score providing a dysfunctional parenting index. Over-reactivity indicates authoritarian and coercive discipline practices; Laxness, in contrast, describes a permissive parent who is inconsistent in providing discipline; Verbosity refers to a parenting style characterized by lengthy and ineffective verbal reprimands. Primary carers were usually mothers and secondary carers (usually fathers).

#### Eye-tracking

The Tobii TX-300 (Tobii AB, Sweden) gaze tracking system was used to record the childrens’ eye movements in 174 infants (58.6%). The battery of tasks comprised a series of animated video clips designed to measure endogenous and exogenous visual attention (Elsabbagh et al., 2013; Elsabbagh et al., 2009; Gliga et al., 2009; Wass et al., 2011). Extracted metrics included baseline orienting speed, visual engagement and disengagement (gap/overlap task), social attention to static and dynamic social scenes and stimuli and cognitive control/reversal learning and visual search. The list of tasks and completion rates are shown in Table 4. All data and extracted metrics are available and can be found at the National Institute for Mental Health (NIMH) data repository portal (https://nda.nih.gov/edit_collection.html?id=3955) (Braithwaite et al., 2023).

**Table 4.** Eye tracking assessment tests and completion rates.

| **Eye-tracking task** | **N (%)** |  |
|---|---|---|
| Gap-overlap | 172 | 57.9% |
| Cognitive control | 170 | 57.2% |
| Non-social contingency | 168 | 56.6% |
| Visual search | 167 | 56.2% |
| Working memory | 167 | 56.2% |
| Fishtanks | 163 | 54.9% |
| Smooth pursuit fixation | 161 | 54.2% |
| Emotions | 159 | 53.5% |
| Fixation | 134 | 45.1% |
| Scenes | 132 | 44.4% |
| Static images | 131 | 44.1% |
| Eye-tracking battery completed | 121 | 40.7% |

### Genomic Data

#### Genetic data

Saliva samples were collected at the initial neonatal MRI data acquisition and 18-month-old infant timepoints using Oragene DNA OG-250 kits (DNAGenotek Inc., Kanata, Canada). The genotyping was performed on only one sample (usually the first). There are no linked maternal or paternal samples. Samples were genotyped on the Illumina Infinium Omni5-4 array v1.2, which comprises a total of 4327108 SNPs, by NIHR BioResource Centre Maudsley Genomics & Biomarker Core Facility. Genotyping was undertaken in two batches. Basic quality control was performed by the Department of Biostatistics & Health Informatics, King’s College London for the combined dHCP batches and a small additional independent study cohort. Raw Illumina microarray genotype image (IDAT) files were uploaded into GenomeStudio and processed according to the GenomeStudio quality control Standard Operating Procedure (https://khp-informatics.github.io/COPILOT/index.html). Data was then further processed according to an in-house pipeline which identified and removed samples with call rates below 95%. It also identified sex mismatches and potential heterozygosity outliers which are flagged in the metadata files. SNP data are available for 154 participants.

#### Methylation Data

Saliva-derived DNA from each sample was treated with sodium bisulfite (Zymo Research EZ-96 DNA Methylation Kit (D5004)). DNA methylation was quantified using the Illumina Infinium HumanMethylationEPIC BeadChip Kit. Methylation analysis for the dHCP samples was undertaken alongside two additional independent study cohorts. A randomized sample layout was generated using key study parameters including all study cohorts, with Omixer R/Bioconductor package (http://www.bioconductor.org/packages/release/bioc/html/Omixer.html). Methylation data are available for 157 participants.

## Discussion

The Developing Human Connectome Project (dHCP) fetal data release sits alongside the corresponding neonatal data release (Edwards et al., 2022) with the intention that they can be useful individually and together to provide marked new insight into the critically important early stages of brain development leading up to and around the time of birth. To maximise comparability between the data from these two collections, all imaging was performed on the same scanner and active steps were taken to harmonise acquisition methods and parameters as far as possible, given the very different constraints that apply to studies in utero, where the mother occupies most of the physical space in the scanner; and in the newborn, when RF coils, patient handling and imaging field of view can all be focused purely on the one subject. Data is released from all steps in the project pipelines, from the native reconstructed images for all modalities, and calibration scans, through intermediate steps in processing, to results from running our processing pipelines. The aim is to allow researchers to work with the data as they wish, without pre-filtering the available selection. In the majority of cases, high-quality images across all modalities are available and are linked to rich collateral data, although practical issues, notably the COVID-19 pandemic, led to some incomplete ascertainment.

The potential of the data has already been highlighted in the first publications using the data to explore different facets of fetal brain development, which together have highlighted how much complexity already exists in the human brain even from 20 weeks gestation, and how the data can provide new insight into the dramatic changes occurring across the period leading up to birth. This has included use of the anatomical data to explore the spatial coupling between gene upregulation during mid-gestation and cortical expansion (Ball et al., 2024) and sulcal morphology (Snyder et al., 2024). The diffusion data has enabled studies delineating the emerging major white matter pathways and their microstructure (Wilson et al., 2021), detailed characterization of the spatiotemporal maturation of the developing thalamocortical pathways and the microstructure of the transient fetal tissue compartments including the subplate and intermediate zone (Wilson et al., 2023), and exploration of the relationship between tissue microstructural maturation and the onset of cortical folding (Wilson et al., 2024). Finally, the functional MRI data has been used to develop novel modelling methods for functional networks which incorporate age-related changes in connectivity directly into network estimation (Karolis et al., 2023).

The data from the dHCP is part of a broader collection of connectome-focused datasets curated within the National Institutes of Mental Health’s extensive multimodal neuroinformatics repository. In combination with the dHCP neonatal data release, this collection of data from fetuses offers a distinctive observational resource, capturing critical insights into the developing human brain during a period of rapid growth and transformation. In addition to the primary MRI data and derived data, additional, complementary, genetic data, follow-up behavioral resources, and atlases are available, providing the most comprehensive resource we could achieve to enable diverse scientific and clinical inquiries. The data are already being widely used, with a selected list of publications available at: https://pubmed.ncbi.nlm.nih.gov/?term=%22developing+human+connectome+project%22&sort=date

## Decleration of the use of AI

We did not use AI in writing this paper.

## ACKNOWLEDGMENTS

We are grateful for all the families who kindly agreed to participate in the project and recognise their particular commitment in remaining engaged with the programme during the COVID-19 Pandemic. We also acknowledge the support of the Newborn Imaging Centre at Evelina London Children’s Hospital.

The Developing Human Connectome Project was funded by the European Research Council under the European Union Seventh Framework Programme (FP/20072013)/ERC Grant Agreement no. 319456. The work was supported by the NIHR Biomedical Research Centres at Guys and St Thomas NHS Trust and the South London and Maudsley NHS Trust, the ESPRC/Wellcome Centre for Medical Engineering and the MRC Centre for Neurodevelopmental Disorders. The views expressed are those of the author(s) and not necessarily those of the NIHR or the Department of Health and Social Care.

TA and VK were supported by a MRC Clinician Scientist Fellowship [MR/P008712/1] and MRC transition support award [MR/V036874/1]. JOM is supported by a Sir Henry Dale Fellowship jointly funded by the Wellcome Trust and the Royal Society [Grant Number 206675/Z/17/Z]. ADE, SW, JOM, TA, and MAR were supported by the Medical Research Council Centre for Neurodevelopmental Disorders [MR/N0266063/1].

LM has been supported by the EU-AIMS (European Autism Interventions) and AIMS-2-TRIALS programmes which receive support from Innovative Medicines Initiative Joint Undertaking Grant No. 115300 and 777394, the resources of which are composed of financial contributions from the European Union’s FP7 and Horizon2020 Programmes, and from the European Federation of Pharmaceutical Industries and Associations (EFPIA) companies’ in-kind contributions, and AUTISM SPEAKS, Autistica and SFARI).

1 Local power scaling, enabled by a direct software modification, made use of measured B1+ field in the fetus.

## REFERENCES

Achenbach, T. M., & Rescorla, L. A. (2000). Manual for the ASEBA Preschool Forms & Profiles.. University of Vermont, Research Center for Children, Youth, & Families.

Achenbach, T. M., & Ruffle, T. M. (2000). The Child Behavior Checklist and related forms for assessing behavioral/emotional problems and competencies. Pediatr Rev, 21(8), 265–271.

Albers, C. A., & Grieve, A. J. (2007). Test Review: Bayley, N. (2006). Bayley Scales of Infant and Toddler Development– Third Edition. San Antonio, TX: Harcourt Assessment. Journal of Psychoeducational Assessment, 25(2), 180–190.

Alexander-Bloch, A., Giedd, J. N., & Bullmore, E. (2013). Imaging structural co-variance between human brain regions. Nat Rev Neurosci, 14(5), 322–336.

Allison, C., Baron-Cohen, S., Wheelwright, S., Charman, T., Richler, J., & Pasco, G. (2008). The Q-CHAT (Quantitative CHecklist for Autism in Toddlers): a normally distributed quantitative measure of autistic traits at 18-24 months of age: preliminary report. J Autism Dev Disord, 38(8), 1414–1425.

Allison, C., Baron-Cohen, S., Wheelwright, S., Charman, T., Richler, J., Pasco, G., & Brayne, C. (2008). The Q-CHAT (Quantitative CHecklist for Autism in Toddlers): a normally distributed quantitative measure of autistic traits at 18-24 months of age: preliminary report. J Autism Dev Disord, 38(8), 1414–1425. 10.1007/s10803-007-0509-7

Arnold, D. S., O’Leary, S. G., Wolff, L. S., & Acker, M. M. (1993). The Parenting Scale: A measure of dysfunctional parenting in discipline situations. Psychological Assessment, 5(2), 137– 144.

Arnold, D. S., O’Leary, S. G., Wolff, L. S., & Acker, M. M. (1993). The Parenting Scale: A measure of dysfunctional parenting in discipline situations.. Psychological Assessment, 5. doi:10.1037/1040-3590.5.2.137

Ball, G., Oldham, S., Kyriakopoulou, V., Williams, L. Z. J., Karolis, V., Price, A., Hutter, J., Seal, M. L., Alexander-Bloch, A., Hajnal, J. V., Edwards, A. D., Robinson, E. C., & Seidlitz, J. (2024). Molecular signatures of cortical expansion in the human foetal brain. Nat Commun, 15(1), 9685. 10.1038/s41467-024-54034-2

Batalle, D., Edwards, A. D., & O’Muircheartaigh, J. (2018). Annual Research Review: Not just a small adult brain: understanding later neurodevelopment through imaging the neonatal brain. J Child Psychol Psychiatry, 59(4), 350–371. 10.1111/jcpp.12838

Bhattacharya, S., Price, A. N., Uus, A., Sousa, H. S., Marenzana, M., Colford, K., Murkin, P., Lee, M., Cordero-Grande, L., Teixeira, R., Malik, S. J., & Deprez, M. (2024). In vivo T2 measurements of the fetal brain using single-shot fast spin echo sequences. Magn Reson Med, 92(2), 715–729. 10.1002/mrm.30094

Bonthrone, A. F., Chew, A., Kelly, C. J., Almedom, L., Simpson, J., & Victor, S. (2021). Cognitive function in toddlers with congenital heart disease: The impact of a stimulating home environment. Infancy, 26(1), 184–199.

Bozek, J., Makropoulos, A., Schuh, A., Fitzgibbon, S., Wright, R., & Glasser, M. F. (2018). Construction of a neonatal cortical surface atlas using Multimodal Surface Matching in the Developing Human Connectome Project. Neuroimage, 179, 11–29.

Bozek, J., Makropoulos, A., Schuh, A., Fitzgibbon, S., Wright, R., Glasser, M. F., Coalson, T. S., O’Muircheartaigh, J., Hutter, J., Price, A. N., Cordero-Grande, L., Teixeira, R., Hughes, E., Tusor, N., Baruteau, K. P., Rutherford, M. A., Edwards, A. D., Hajnal, J. V., Smith, S. M.,…Robinson, E. C. (2018). Construction of a neonatal cortical surface atlas using Multimodal Surface Matching in the Developing Human Connectome Project. Neuroimage, 179, 11–29. 10.1016/j.neuroimage.2018.06.018

Braithwaite, E., Kyriakopoulou, V., Mason, L., Davidson, A., Tusor, N., & Harper, N. (2023). Objective assessment of visual attention in toddlerhood [Preprint].

Christiaens, D., Cordero-Grande, L., Pietsch, M., Hutter, J., Price, A. N., & Hughes, E. J. (2021). Scattered slice SHARD reconstruction for motion correction in multi-shell diffusion MRI. Neuroimage, 225, 117437.

Christiaens, D., Cordero Grande, L., Price, A., Hutter, J., Hughes, E., Counsell, S., Tournier, J. D., & Hajnal, J. V. (2019). Fetal diffusion MRI acquisition and analysis in the developing Human Connectome Project. Proceedings of Int Soc Magn Reson Med, Montreal.

Christiaens, D., Pietsch, M., Cordero-Grande, L., Price, A. N., Hutter, J., Hughes, E. J., Counsell, S. J., Hajnal, J. V., & Tournier, J. D. (2021). Combined spin echo and gradient echo slice- to-volume reconstruction in fetal diffusion MRI. Proceedings of Int Soc Magn Reson Med,

Cordero-Grande, L., Christiaens, D., Hutter, J., Price, A. N., & Hajnal, J. V. (2019). Complex diffusion-weighted image estimation via matrix recovery under general noise models. NeuroImage, 200(December 2018), 391–404.

Cordero-Grande, L., Price, A., Ferrazzi, G., Hutter, J., Christiaens, D., Hughes, E. J., & Hajnal, J. V. (2018). Spin And Field Echo (SAFE) dynamic field correction in 3T fetal EPI. Proceedings of Int Soc Magn Reson Med,

Cordero-grande, L., Price, A. N., Hughes, E. J., Wright, R., Rutherford, M. A., & Hajnal, J. V. (2019). Automating fetal brain reconstruction using distance regression learning. Proceedings of Int Soc Magn Reson Med,

Cosío-Guirado, R., Tapia-Medina, M. G., Kaya, C., Peró-Cebollero, M., Villuendas-González, E. R., & Guàrdia-Olmos, J. (2024). A comprehensive systematic review of fMRI studies on brain connectivity in healthy children and adolescents: Current insights and future directions. Developmental Cognitive Neuroscience, 69, 101438. 10.1016/j.dcn.2024.101438

Craddock, R. C., Jbabdi, S., Yan, C. G., Vogelstein, J. T., Castellanos, F. X., & Di Martino, A. (2013). Imaging human connectomes at the macroscale. Nat Methods, 10(6), 524–539.

Edwards, A. D., Rueckert, D., Smith, S. M., Abo Seada, S., Alansary, A., & Almalbis, J. (2022). The Developing Human Connectome Project Neonatal Data Release. Front Neurosci, 16, 886772.

Elsabbagh, M., Gliga, T., Pickles, A., Hudry, K., Charman, T., & Johnson, M. H. (2013). The development of face orienting mechanisms in infants at-risk for autism. Behav Brain Res, 251, 147–154.

Elsabbagh, M., Volein, A., Holmboe, K., Tucker, L., Csibra, G., & Baron-Cohen, S. (2009). Visual orienting in the early broader autism phenotype: disengagement and facilitation. J Child Psychol Psychiatry, 50(5), 637–642.

Ferrazzi, G. (2016). An exploration of methods for performing resting state fMRI in the human fetus King’s College London]. London.

Ferrazzi, G., Price, A. N., Teixeira, R., Cordero-Grande, L., Hutter, J., & Gomes, A. (2018). An efficient sequence for fetal brain imaging at 3T with enhanced T(1) contrast and motion robustness. Magn Reson Med, 80(1), 137–146.

Fischl, B. (2012). FreeSurfer. Neuroimage, 62(2), 774–781. 10.1016/j.neuroimage.2012.01.021

Gaspar, A., Nunes, R. G., Ferrazzi, G., Hughes, E. J., Hutter, J., & Malik, S. J. (2018). Optimizing maternal fat suppression with constrained image-based shimming in fetal MR. Magn Reson Med, 81(1), 477–485.

Gliga, T., Elsabbagh, M., Andravizou, A., & Johnson, M. (2009). Faces Attract Infants’ Attention in Complex Displays. Infancy, 14(5), 550–562. 10.1080/15250000903144199

Haataja, L., Mercuri, E., Regev, R., Cowan, F., Rutherford, M., & Dubowitz, V. (1999). Optimality score for the neurologic examination of the infant at 12 and 18 months of age. J Pediatr, *135*(2 Pt 1), 153–161.

Hisle-Gorman, E., Susi, A., Stokes, T., Gorman, G., Erdie-Lalena, C., & Nylund, C. M. (2018). Prenatal, perinatal, and neonatal risk factors of autism spectrum disorder. Pediatr Res, 84(2), 190–198. 10.1038/pr.2018.23

Hughes, E. J., Price, A. N., McCabe, L., Hiscocks, S., Waite, L., & Green, E. (2021). The effect of maternal position on venous return for pregnant women during MRI. NMR Biomed, 34(4), e4475.

Hutter, J., Price, A. N., Cordero-Grande, L., Malik, S., Ferrazzi, G., Gaspar, A., Hughes, E. J., Christiaens, D., McCabe, L., Schneider, T., Rutherford, M. A., & Hajnal, J. V. (2018). Quiet echo planar imaging for functional and diffusion MRI. Magn Reson Med, 79(3), 1447–1459. 10.1002/mrm.26810

Hutter, J., Tournier, J. D., Price, A. N., Cordero-Grande, L., Hughes, E. J., & Malik, S. (2018). Time-efficient and flexible design of optimized multishell HARDI diffusion. Magn Reson Med, 79(3), 1276–1292.

Jeurissen, B., Tournier, J. D., Dhollander, T., Connelly, A., & Sijbers, J. (2014). Multi-tissue constrained spherical deconvolution for improved analysis of multi-shell diffusion MRI data. Neuroimage, 103, 411–426. 10.1016/j.neuroimage.2014.07.061

Karolis, V. R., Cordero-Grande, L., Price, A. N., Hughes, E., Fitzgibbon, S. P., Kyriakopoulou, V., Uus, A., Harper, N., Prokopenko, D., Bridglal, D., Moore, J. W., Wilson, S., Pietsch, M., Christiaens, D., Deprez, M., Williams, L. Z. J., Robinson, E. C., Makropoulos, A., Farahibozorg, S. R.,…Hajnal, J. V. (2025). The developing Human Connectome Project fetal functional MRI release: Methods and data structures. Imaging Neurosci (Camb), 3. 10.1162/imag_a_00512

Karolis, V. R., Fitzgibbon, S. P., Cordero-Grande, L., Farahibozorg, S. R., Price, A. N., & Hughes, E. J. (2023). Maturational networks of human fetal brain activity reveal emerging connectivity patterns prior to ex-utero exposure. Commun Biol, 6(1), 661.

Kuklisova-Murgasova, M., Quaghebeur, G., Rutherford, M. A., Hajnal, J. V., & Schnabel, J. A. (2012). Reconstruction of fetal brain MRI with intensity matching and complete outlier removal. Medical Image Analysis, 16(8), 1550–1564.

Liang, K., Guo, Y., Williams, L. Z. J., Besenczi, R., Sun, Z., David Edwards, A., Robinson, E. C., & Nosarti, C. (2025). A cortical signature of very preterm birth across development and its association with neurodevelopmental outcomes. BioRxiv, 2025.2010.2017.683086. 10.1101/2025.10.17.683086

Ma, Q., Kyriakopoulou, V., Hajnal, J., Robinson, E., Kainz, B., & Rueckert, D. (2023). Conditional temporal attention networks for neonatal cortical surface reconstruction. International Conference on Medical Image Computing and Computer-Assisted Intervention.,

Makropoulos, A., Robinson, E. C., Schuh, A., Wright, R., Fitzgibbon, S., & Bozek, J. (2018). The developing human connectome project: A minimal processing pipeline for neonatal cortical surface reconstruction. Neuroimage, 173, 88–112.

Molnár, Z., Clowry, G. J., Šestan, N., Alzu’bi, A., Bakken, T., Hevner, R. F., Hüppi, P. S., Kostović, I., Rakic, P., Anton, E. S., Edwards, D., Garcez, P., Hoerder-Suabedissen, A., & Kriegstein, A. (2019). New insights into the development of the human cerebral cortex. J Anat, 235(3), 432–451. 10.1111/joa.13055

Montagna, A., & Nosarti, C. (2016). Socio-Emotional Development Following Very Preterm Birth: Pathways to Psychopathology. Front Psychol, 7, 80. 10.3389/fpsyg.2016.00080

Nehrke, K., & Bornert, P. (2012). DREAM--a novel approach for robust, ultrafast, multislice B(1) mapping. Magn Reson Med, 68(5), 1517–1526.

Pietsch, M., Christiaens, D., Hutter, J., Cordero-Grande, L., Price, A. N., & Hughes, E. (2019). A framework for multi-component analysis of diffusion MRI data over the neonatal period. Neuroimage, 186, 321–337.

Price, A. N., Cordero-Grande, L., Hughes, E. J., Hiscocks, S., Green, E., McCabe, L., Hutter, J., Ferrazzi, G., Deprez, M., Roberts, T., Christiaens, D., Duff, E., Karolis, V., Malik, S. J., Rutherford, M. A., Edwards, A. D., & Hajnal, J. V. (2019). The developing Human Connectome Project ( dHCP ): fetal acquisition protocol. Proceedings of Int Soc Magn Reson Med,

Price, A. N., Cordero-Grande, L., Malik, S. J., & Hajnal, J. V. (2018). Multiband zoom TSE imaging: increasing efficiency with multiband tip-back preparation pulses. Proceedings of Int Soc Magn Reson Med, Paris.

Putnam, S. P., Gartstein, M. A., & Rothbart, M. K. (2006). Measurement of fine-grained aspects of toddler temperament: the early childhood behavior questionnaire. Infant Behav Dev, 29(3), 386–401. 10.1016/j.infbeh.2006.01.004

Putnam, S. P., & Rothbart, M. K. (2006). Development of short and very short forms of the Children’s Behavior Questionnaire. J Pers Assess, 87(1), 102–112.

Robinson, E. C., Garcia, K., Glasser, M. F., Chen, Z., Coalson, T. S., Makropoulos, A., Bozek, J., Wright, R., Schuh, A., Webster, M., Hutter, J., Price, A., Cordero Grande, L., Hughes, E., Tusor, N., Bayly, P. V., Van Essen, D. C., Smith, S. M., Edwards, A. D.,…Rueckert, D. (2018). Multimodal surface matching with higher-order smoothness constraints. Neuroimage, 167, 453–465. 10.1016/j.neuroimage.2017.10.037

Rutherford, M. A. (2009). Magnetic resonance imaging of the fetal brain. Curr Opin Obstet Gynecol, 21(2), 180–186. 10.1097/GCO.0b013e32832947ab

Schuh, A., Makropoulos, A., Robinson, E. C., Cordero-Grande, L., Hughes, E., & Hutter, J. (2018). Unbiased construction of a temporally consistent morphological atlas of neonatal brain development [Preprint].

Schuh, A., Murgasova, M., Makropoulos, A., Ledig, C., Counsell, S. J., Hajnal, J. V., Aljabar, P., & Rueckert, D. (2014). Construction of a 4D Brain Atlas and Growth Model Using Diffeomorphic Registration. MICCAI STIA,

Serag, A., Aljabar, P., Ball, G., Counsell, S. J., Boardman, J. P., Rutherford, M. A., Edwards, A. D., Hajnal, J. V., & Rueckert, D. (2012). Construction of a consistent high-definition spatio-temporal atlas of the developing brain using adaptive kernel regression. Neuroimage, 59(3), 2255–2265. S1053-8119(11)01136-0 [pii] 10.1016/j.neuroimage.2011.09.062

Smith, S. M., Jenkinson, M., Woolrich, M. W., Beckmann, C. F., Behrens, T. E., & Johansen-Berg, H. (2004). Advances in functional and structural MR image analysis and implementation as FSL. Neuroimage, 23(Suppl 1), S208–S219.

Snyder, W. E., Vertes, P. E., Kyriakopoulou, V., Wagstyl, K., Williams, L. Z. J., Moraczewski, D., Thomas, A. G., Karolis, V. R., Seidlitz, J., Riviere, D., Robinson, E. C., Mangin, J. F., Raznahan, A., & Bullmore, E. T. (2024). A bimodal taxonomy of adult human brain sulcal morphology related to timing of fetal sulcation and trans-sulcal gene expression gradients. Neuron, 112(20), 3396–3411 e3396. 10.1016/j.neuron.2024.07.023

Tournier, J. D., Christiaens, D., Hutter, J., Price, A. N., Cordero-Grande, L., & Hughes, E. (2020). A data-driven approach to optimising the encoding for multi-shell diffusion MRI with application to neonatal imaging. NMR Biomed, 33(9), e4348.

Tournier, J. D., Smith, R. E., Raffelt, D., Tabbara, R., Dhollander, T., & Pietsch, M. (2019). MRtrix3: A fast, flexible and open software framework for medical image processing and visualisation. Neuroimage, 202, 116–137.

Tustison, N. J., Cook, P. A., Holbrook, A. J., Johnson, H. J., Muschelli, J., Devenyi, G. A., Duda, J. T., Das, S. R., Cullen, N. C., Gillen, D. L., Yassa, M. A., Stone, J. R., Gee, J. C., & Avants, B. B. (2021). The ANTsX ecosystem for quantitative biological and medical imaging. Sci Rep, 11(1), 9068. 10.1038/s41598-021-87564-6

Uus, A., Fukami-Gartner, A., Kyriakopoulou, V., Cromb, D., Morgan, T., Arulkumaran, S., Collado, A. E., Luis, A., Bos, R., Makropoulos, A., Schuh, A., Robinson, E., Sousa, H., Deprez, M., Cordero-Grande, L., Bradshaw, C., Colford, K., Hutter, J., Price, A.,…Story, L. (2026). Multi-BOUNTI: Multi-lobe Brain vOlUmetry and segmeNtation for feTal and neonatal MRI. medRxiv, 2026.2004.2021.26351376. 10.64898/2026.04.21.26351376

Uus, A., Hall, M., Payette, K., Hajnal, J. V., Deprez, M., Rutherford, M. A., Hutter, J., & Story, L. (2023). Combined Quantitative T2* Map and Structural T2-Weighted Tissue-Specific Analysis for Fetal Brain MRI: Pilot Automated Pipeline. Perinatal, Preterm and Paediatric Image Analysis, Cham.

Uus, A., Kyriakopoulou, V., Cordero-Grande, L., Christiaens, D., Pietsch, M., & Price, A. (2023). Multi-channel spatio-temporal MRI atlas of the normal fetal brain development from the developing Human Connectome Project. In: G-Node Open Data.

Van Essen, D. C., Smith, S. M., Barch, D. M., Behrens, T. E., Yacoub, E., & Ugurbil, K. (2013). The WU-Minn Human Connectome Project: an overview. Neuroimage, 80, 62–79.

Wass, S., Porayska-Pomsta, K., & Johnson, M. H. (2011). Training attentional control in infancy. Curr Biol, 21(18), 1543–1547.

Williams, L. Z. J., Fitzgibbon, S. P., Bozek, J., Winkler, A. M., Dimitrova, R., Poppe, T., Schuh, A., Makropoulos, A., Cupitt, J., O’Muircheartaigh, J., Duff, E. P., Cordero-Grande, L., Price, A. N., Hajnal, J. V., Rueckert, D., Smith, S. M., Edwards, A. D., & Robinson, E. C. (2023). Structural and functional asymmetry of the neonatal cerebral cortex. Nat Hum Behav, 7(6), 942–955. 10.1038/s41562-023-01542-8

Wilson, S., Christiaens, D., Yun, H., Uus, A., Cordero-Grande, L., Karolis, V., Price, A., Deprez, M., Tournier, J. D., Rutherford, M., Grant, E., Hajnal, J. V., Edwards, A. D., Arichi, T., O’Muircheartaigh, J., & Im, K. (2024). Dynamic changes in subplate and cortical plate microstructure at the onset of cortical folding in vivo. BioRxiv. 10.1101/2023.10.16.562524

Wilson, S., Pietsch, M., Cordero-Grande, L., Christiaens, D., Uus, A., Karolis, V. R., Kyriakopoulou, V., Colford, K., Price, A. N., Hutter, J., Rutherford, M. A., Hughes, E. J., Counsell, S. J., Tournier, J. D., Hajnal, J. V., Edwards, A. D., O’Muircheartaigh, J., & Arichi, T. (2023). Spatiotemporal tissue maturation of thalamocortical pathways in the human fetal brain. Elife, 12. 10.7554/eLife.83727

Wilson, S., Pietsch, M., Cordero-Grande, L., Price, A. N., Hutter, J., & Xiao, J. (2021). Development of human white matter pathways in utero over the second and third trimester. Proceedings of the National Academy of Sciences, 118(20).

Wolke, D., Jaekel, J., Hall, J., & Baumann, N. (2013). Effects of sensitive parenting on the academic resilience of very preterm and very low birth weight adolescents. J Adolesc Health, 53(5), 642–647.

Wright, R., Khanal, B., Gomez, A., Skelton, E., Matthew, J., Hajnal, J. V., Rueckert, D., & Schnabel, J. A. (2018). LSTM Spatial Co-transformer Networks for Registration of 3D Fetal US and MR Brain Images. MICCAI,

Zalesky, A., & Di Biase, M. A. (2026). Clinical connectomics: Mapping structural brain networks in disease. Neuron, 114(12), 2089–2108. 10.1016/j.neuron.2026.02.007

Zhao, Z., Zhang, H., Ma, C., Fan, C., & Zhao, H. (2020). Comparative study of phase unwrapping algorithms based on solving the Poisson equation. Measurement Science and Technology, 31(6).

